# Undergraduate student interest in healthcare career in the context of COVID-19 pandemic

**DOI:** 10.1101/2021.04.11.438530

**Authors:** Stephanie Y Jo, Jingxin Li, Ananya Dewan, Yu-Chia Cheng, Haoxiang Hou, Ronnie A. Sebro

## Abstract

**Objectives:** The healthcare profession has been long considered an excellent career choice. Pre-medical experience is documented to be important in shaping future medical landscape. In the wake of the pandemic, there has been intense media spotlight on the healthcare profession and change in academic environment, necessitating analyses of student experience. This project aims to assess change in undergraduate student interest in healthcare career using cross-sectional survey study.

**Methods:** The project was approved by our Institutional Review Board. Voluntary survey collected data on demographics, socioeconomics, media exposure, academic environment, and change in interest in a healthcare profession. Survey was distributed through the university undergraduate pre-health listserv. Total of 297 responses were obtained. Descriptive statistics including Fisher’s exact test were applied in the analysis.

**Results:** Majority of the respondents were Asians (54.9%), second generation immigrants (52.2%), and female (73.4%). Large proportion of the respondents were negatively affected by the pandemic, with losing a job or internship personally (42.1%) or a family member or a friend (62.6%). Students had mixed response to online learning environment, with 27.3% of students noting no change, 40.4% students noting increased difficulty, and 32.3% students noting decreased difficulty of classes. During the pandemic, 47.5% of students noted increased interest in pursuing healthcare career. The change in interest in healthcare career was not associated with xdemographics, economic hardship, or online learning environment.

**Discussion:** Despite the challenges of COVID-19 pandemic, students showed strong interest in pursuing healthcare careers.

## Introduction

The healthcare profession has long been considered an excellent career choice with fulfillment of desire to help others (1), possibility of high income (2) and job security (3). However, surveys of the general public demonstrate that there is also a substantial level of mistrust in medical professionals (4). Moreover, physician burnout is a crisis that is becoming increasing recognized as more and more physicians are feeling overworked and undervalued (5). During the already trying times, SARS-CoV-2 virus hit the world with global pandemic. Given the enormous effect the pandemic had on everyday lives, it is imperative we understand the influence of the decision to pursue a career in healthcare by students.

Previous study shows that pre-medical experience in undergraduate years is important in attrition from premedical track and well-being of physicians (1). Hence, with the extensive effect of COVID-19 pandemic strongly affecting the social interaction, economy, and learning environment, there is need for analysis of undergraduate students’ experience. In particular, the majority of educational experience has shifted online, and there has been intense media spotlight on the healthcare profession. It is unknown how disruptions to academic environment and the broadcasting of dangers and stress of healthcare professional affect a student’s choice to enter healthcare career. Although similar studies have been conducted with medical students (6), one has not been conducted with undergraduate students. This project surveyed demographic, socioeconomic, and academic factors that may affect an undergraduate student’s interest in healthcare profession and aims to assess change in interest during the pandemic. This information will give insight into characteristics of future workforce in healthcare profession.

## Methods

Study was approved by the institutional review board. After the approval, the survey was distributed through the pre-health listserv to the undergraduate students through self-administered electronic link. Students interested in nursing school or medical school can voluntarily sign up for subscription to the pre-health listserv. Cross-sectional online survey response was obtained from July 22, 2020 to November 17, 2020. The survey questions were designed by authors and consisted of demographic and socioeconomic information, interest in healthcare career, academic environment, and time spent watching, listening, or reading news (media exposure). Complete survey is attached as supplementary material. As incentive to take the survey, survey respondents had the option to leave personal email address for entry into a $5 Amazon gift card raffle. Otherwise, survey participation was voluntary and no identifying information was obtained. Survey responses were collected through Google survey application.

The survey link email was opened by 648 recipients. Total of 297 responses were obtained with response rate of 45.8% (297/648). Two-sided Fisher’s exact test used to compare proportions. P-values less than 0.05 were considered statistically significant. Data were analyzed utilizing R software version 3.6.

## Results

Survey link was distributed to pre-health listserv at our institution. Pre-health listserv subscribers include students interested in pursuing medical school and nursing school and was considered appropriate survey group. Demographic results of survey respondents were as follows. Age of the respondents ranged from 17 to 31, with 95.3% (283/297) of respondents between ages 18 and 23. 73.4% (218/297) identified as female. Students’ hometowns were from all across the United States as defined by the U.S. Census Bureau (7); majority 46.5% (138/297) were from Northeast and 23.9% (71/297) were from South. This information was obtained as different parts of the country were differentially affected early in the pandemic. According to the federal government racial and ethnic categories (8), majority 54.9% (163/297) were Asians and 26.9% (80/297) were White. Others were Hispanic or Latino, African Americans, or multiracial. This information was obtained as minority groups are disproportionately affected by the pandemic. Majority of the students 52.2% (155/297) were second generation immigrants (student naturally had United States citizenship from birth, but not so for parents). Next large group was those who had been in the United States for over three generations (21.5%, 64/297). 11.4% (34/297) were first generation immigrants (student did not have United States citizenship by birth but obtained one), and 6.4% (19/297) were third generation immigrant (student and at least one parent naturally had USA citizenship from birth, but not so for grandparents). Rest 2.7% were international students, adopted, don’t know, or choose not to answer. 58.2% (173/297) student were fluent in language other than English that was used at home. These results show that the pre-health listserv included high proportion of women, minorities, and immigrant families.

Next, we evaluated the economic aspect of our students. 24.9% (74/297) students relied solely on parental support for undergraduate tuition payment. Others used combination of parental support, financial aid, work study, scholarship, and work outside of school to cover tuition. 73.4% (218/297) of students were dependent financially. 23.9% (71/297) of students supported themselves, and 1.3% (4/297) of students supported other family members. Pandemic had strong negative economic impact on the student community, with 42.1% (125/297) of students reporting losing a job or an internship position due to the pandemic personally. Larger proportion (62.6%, 186/297) of students had a family member or a friend lose a job or an internship position due to the pandemic. This information was obtained as economic hardship may impact a student’s interest in pursuing additional schooling.

We then evaluated the academic environment of the students. Survey respondents were distributed throughout the undergraduate years; 27.3 % (81/297) students were first year in college, 27.3% (81/297) students were second year in college, 16.2% (48/297) students were third year in college, 17.8% (53/297) students were fourth year in college. This question was asked as students further along the undergraduate studies may be more fixed in career path. With the onset of pandemic and state-wide lockdown in spring of 2020, our institution switched to online learning. Majority (76.1%, 226/297) of students had combination of live lecture session and recorded lecture session. This was accompanied by additional help through emails and/or live office hours. 27.3% (81/297) of students thought the difficulty of classes did not change with the socially distance classroom. 32.3% (96/297) of students thought the classes easier and 40.4% (120/297) of students thought the classes were harder. The perceived class difficulty may change pursuit of professional school, with those feeling the classes got easier may be encouraged and those feeling the classes got harder discouraged.

There has been frequent coverage of coronavirus pandemic in the media. High exposure to the pandemic news may affect a student’s decision to pursuit healthcare career. Time spent watching, reading, or listening to news (media exposure) per day and source of these news were surveyed. 42.1% (125/297) students had up to 1 hour of media exposure, and 6.7% (20/297) students had little to no exposure to media. 35.4% (105/297) had 1-2 hours of media exposure, 11.1% (33/297) students had 2-4 hours of media exposure, and 4.0% (12/297) students had more than 4 hours of media exposure per day. 9.1% (27/297) relied on family, friends, and word of mouth for news, 11.8% (35/297) relied on federal, state, and local government websites, and majority relied on national news outlet (e.g., CNN, NBC) 52.2% (155/297). 24.9% (74/297) relied on social media for news. Others noted mixture of news source. Survey responses are summarized in Table 1.

**Table 1.** Summary of survey responses.

|  |  |
| --- | --- |
| Gender Identity |  |
| Female | 73.4% (218/297) |
| Male | 25.9% (77/297) |
| Other | 0.7% (2/297) |
| Race/ Ethnicity |  |
| Asian | 54.9% (163/297) |
| White | 26.9% (80/297) |
| Hispanic or Latino | 6.4% (19/297) |
| Black | 5.7% (17/297) |
| Multi-racial | 5.4% (16/297) |
| Other | 0.7% (2/297) |
| Hometown |  |
| Northeast | 46.5% (138/297) |
| South | 23.9% (71/297) |
| Midwest | 10.4% (31/297) |
| West | 10.1% (30/297) |
| International | 8.7% (26/297) |
| Other | 0.3% (1/297) |
| Immigration status |  |
| First generation immigrants (student did not have United States citizenship by birth but obtained one) | 11.4% (34/297) |
| Second generation immigrants (student naturally had United States citizenship from birth, but not so for parents) | 52.2% (155/297) |
| Third generation immigrant (student and at least one parent naturally had USA citizenship from birth, but not so for grandparents) | 6.4% (19/297) |
| 3+ generation in the United States | 21.5% (64/297) |
| Other (international, adopted, don't know) | 7.1% (21/297) |
| Financial independence status |  |
| Dependent | 74.7% (222/297) |
| Self-supported | 23.9% (71/297) |
| Supporting others | 1.3% (4/297) |
| Economic hardship |  |
| Lost job or internship position – self | 42.1% (125/297) |
| Lost job or internship position – family or friend | 62.6% (186/297) |
| Academic environment |  |
| Classes became easier due to online format | 32.3% (96/297) |
| No change in difficulty | 27.3% (81/297) |
| Classes became harder due to online format | 40.4% (120/297) |
| Media exposure |  |
| Little to none | 6.7% (20/297) |
| Up to 1 hour | 42.1% (125/297) |
| 1-2 hours | 35.4% (105/297) |
| 2-4 hours | 11.1% (33/297) |
| 4+ hours | 4.0% (12/297) |
| Interest in healthcare career |  |
| Increased | 47.5% (141/297) |
| Unchanged | 32.3% (96/297) |
| Decreased | 2.7% (8/297) |

We next evaluated if there was change in students’ interest in pursuing healthcare career. 47.5% (141/297) of students noted increase in interest in healthcare career, and 32.3% (96/297) of students noted no change in interest in healthcare career during the pandemic. Only 2.7% (8/297) of students noted decrease in interest in healthcare career. This result shows that pandemic situation heightened students’ interest in entering healthcare career. We also analyzed if the increased interested was associated with other survey responses utilizing two-sided Fisher’s exact test. Surprisingly, the interest in pursuing healthcare career was not affected by geographic location, race/ ethnicity, immigration status, economic hardship, media exposure, or increased or decreased difficulty of classes. Results are summarized in Table 2.

**Table 2.** Fisher’s exact test of association with interest in healthcare career.

|  |  |  |
| --- | --- | --- |
| Geographic location and interest in healthcare post COVID (p = 0.087) |  |  |
|  | Unchanged/decreased | Increased |
| Midwest | 17 | 11 |
| Northeast | 70 | 68 |
| Outside US | 11 | 16 |
| South | 36 | 38 |
| West | 22 | 8 |
| Race/ethnicity and interest in healthcare post COVID (p = 0.2368) |  |  |
|  | Unchanged/decreased | Increased |
| Asian | 85 | 78 |
| Black/African American | 6 | 11 |
| White | 49 | 31 |
| Hispanic or Latino | 9 | 10 |
| Multiracial or multiethnic, other | 7 | 11 |
| Immigration status and interest in healthcare post COVID (p = 0.769) |  |  |
|  | Unchanged/decreased | Increased |
| First generation | 16 | 20 |
| Second generation immigrants | 83 | 72 |
| Third generation immigrants | 10 | 9 |
| USA 3+ generation | 37 | 27 |
| International, other | 8 | 11 |
| Personal economic hardship and interest in healthcare post COVID (p = 0.725) |  |  |
|  | Unchanged/decreased | Increased |
| No | 88 | 83 |
| Yes | 67 | 58 |
| Family/ friend economic hardship and interest in healthcare post COVID (p = 0.4702) |  |  |
|  | Unchanged/decreased | Increased |
| No | 61 | 49 |
| Yes | 94 | 92 |
| Class year and interest in healthcare post COVID (p = 0.853) |  |  |
|  | Unchanged/decreased | Increased |
| Freshmen | 44 | 37 |
| Sophomore | 45 | 36 |
| Junior | 22 | 26 |
| Senior | 27 | 24 |
| Other | 18 | 18 |
| Perceived difficulty of classes and interest in healthcare post COVID (p = 0.103) |  |  |
|  | Unchanged/decreased | Increased |
| Unchanged | 43 | 38 |
| Easier | 58 | 38 |
| Harder | 55 | 65 |
| Media exposure and interest in healthcare post COVID (p = 0.181) |  |  |
|  | Unchanged/decreased | Increased |
| Little to none | 12 | 10 |
| < 1 hr | 75 | 50 |
| 1-2 | 49 | 56 |
| 2-4 | 16 | 17 |
| 4+ hours | 4 | 8 |

## Discussion

While initially affecting limited geographical regions during spring of 2020, our survey responses were obtained when the entire United States and the world became affected by the SARS-Cov-2 virus. This cross-sectional study aimed to evaluate the undergraduate students’ experience during the pandemic and its potential effect on the decision to enter healthcare career. Overall, the interest in pursuing healthcare career has increased or remained unchanged. The factors we examined that may influence career decision, including demographics, economic hardship, media exposure, and online learning environment, did not change students’ interest in healthcare career. This may in part due to the characteristics of survey respondents, who were subscribed to pre-health listserv and already interested in medical or nursing career. It is reassuring that pandemic promoted interest in healthcare career, rather than dissuading students from entering the field.

While we did not find a factor that was associated with change in healthcare career interest, a few interesting observations on future healthcare workforce have emerged from this study. Frist, there were high proportions of Asians (54.9%) and second-generation immigrants (52.2%) in the pre-health listserv. There was also high proportion of students who were fluent in non-English language used at home (58.2%). This points to the increasing diversity in the healthcare workforce. In fact, according to an article from 2016, foreign born doctors made up 25% of all physicians and Asian American medical school graduates made up approximately 20% of the class (9). It is unclear why there is high number of Asians and second-generation immigrants interested in healthcare career. There may be strong cultural preference in the Asian community in pursuing career in healthcare. Also, healthcare is consistently the top ten highest professions to receive visa approval (10), which may facilitate settling in the United States and thus have positive reputation in immigrant communities. It will be interesting to assess pre-health listserv demographics in other institutions to see if the trend is similar. Additionally, it is still imperative that we continue to find ways to encourage students who are traditionally underrepresented in medicine to consider the profession and find ways to support them in their pursuits.

Second, large majority (73.4%) of the respondents were females. There has been increasing female matriculants in medical school. In fact, in 2019, women comprised slight majority 50.5% of all medical school students (11). Large proportion of female respondents in our survey reflects the trend of women workforce in healthcare. With this trend of increasing women and minority as healthcare professionals, there is greater mandate to support our workforce. Multiple studies have shown that there is patient bias against women and minority healthcare workers (12–14), and these biased encounters have negative effects on professional and personal identity. In order to best serve patients and protect the workforce, healthcare as a field must set firm boundaries against bias and aim for embracing diverse and inclusive work environment.

Main limitations of our study are that it is restricted to a single institution, and the racial/ ethnic distribution and immigration status were heavily skewed and not reflective of the general population. Further studies including additional institutions will be helpful to better gauge the demographics of undergraduate students pursuing healthcare careers. Another limitation is that the email link was sent to pre-health listserv, which is a biased student cohort with already high interest in healthcare career. Those who were not interested in healthcare career previously and now plan to pursue healthcare will not be captured by the survey. Also, among the students that opened the email with the survey link, approximately half did not respond. While it is limitation of all survey-based study, the students who were willing to participate in the survey may be different from those who did not respond. On the other hand, among those who were already interested, there was persistent interest despite the pandemic.

There has been large increase in application to medical school in 2020, which has been compared to “time after 9/11, when we saw an increase in those motivated to serve this country militarily” (15) It is great to see the desire to serve the community by our students, and this corresponds to our finding of increased interest in healthcare career. The media spotlight of the pandemic may emphasize the value of the healthcare professional and may spark a trend of increased interest in the field. Multi-year study will be necessary to assess the long-term effects of pandemic on the healthcare profession. As we welcome new members into this profession, we must continue our efforts at supporting and meeting the need of increasingly diverse workforce.

## Supporting information

survey questions

