## Supplementary material for "Undergraduate student interest in healthcare career in the context of COVID-19 pandemic": survey questions

Healthcare Career in COVID-19 Pandemic Survey
https://docs.google.com/forms/d/e/1FAIpQLSd0bcb2SZ9rAz-9xtSgkXGN1SLgRVPr2281IQsJcUw3wvdqgA/viewform

Page 1
Your participation is completely voluntary, and you can withdraw at any time.
To take this survey, you must be:
- At least 18 years old
- Be an undergraduate student, post-baccalaureate student, currently considering or at one time considered becoming a healthcare professional

All data will remain anonymous. If you choose to provide your email to be entered into the drawing for one of several $5 amazon gift cards, your email will not be used in conjunction with the answers you provided.

A detailed consent form and study information can be found here: https://drive.google.com/file/d/1O9DtBbLgQTZHhySXi2H0DqXg_QPdgZ8Q/view?usp=sharing

If you meet these criteria and would like to take the survey, indicate below and continue to start

Page 2
By selecting 'Yes' you are indicating you have read the informed consent and agree to participate in the research study *

Survey
What is your age? *

What is your gender? *
- Female
- Male
- Non Binary
- Other:

What is your race or ethnicity? *
- Native American or Alaska Native
- Caucasian
- Multiracial or Multiethnic
- Asian
- Native Hawaiian or other Pacific Islander
- Black or African American
- Hispanic or Latino
- Middle Eastern or North African
- Other:

What is your hometown (State, Country) *

What is your hometown city? *

What is your Immigration status? *

- First generation immigrants (yourself do not have USA citizenship or initially did not have USA citizenship but obtained one)
- Second generation immigrants (yourself naturally had USA citizenship from birth, but not so for parents)
- Third generation immigrants (yourself and at least one parent naturally had USA citizenship from birth, but not so for grandparents)
- In USA 3+ generation
- International
- I choose not to answer or don't know
- Other:

Fluent language(s) other than English used at home

What College or University do you currently attend? *

How is your tuition paid? *

- Loan (need to be repaid)
- Parental support
- Financial Aid
- Scholarship (does not need to be repaid)
- Work during school (excluding work study)
- Work study related to financial aid
- Other:

Are you taking out an educational loan? (Please specify amount that needs to be paid back)

- none
- < $10,000
- $10,000 - $25,000
- $25,000 - $50,000
- $50,000 - $100,000
- $100,000 - $150,000
- $150,000 - $300,000
- +$300,000

Are you supporting anyone financially? *

- None
- Self
- Child / Children
- Parents / Extended family
- Spouse / Partner

- Other:

Year in college for Fall 2020 *
- Undergraduate year 1
- Undergraduate year 2
- Undergraduate year 3
- Undergraduate year 4
- Undergraduate year 4+
- Taking semester / year off of school
- Post-Bac year
- Other:

What is/was your major / minor in college? *Select all that apply
- Undecided
- Arts and Humanities
- Business
- Pre-Med
- Multi-/Interdisciplinary Studies
- Public and Social Services
- Science, Math, and Technology
- Social Sciences
- Trades and Personal Services
- Other:
- What career field are you planning to pursue after college? *
- Undecided
- Education
- Finance/ Business
- Law
- Administration
- IT/Tech/Engineering
- Physician/Medicine
- Other Healthcare Professional (e.g. nursing, physician assistant, pharmacist, dentist, etc)
- Other:

What kind of healthcare professional are you working towards becoming? *
- Nurse/ Nurse Practitioner/ Nurse Anesthetist
- Physical/ occupational/ Speech Therapist
- Pharmacist
- Medical technician/ Dental hygienist
- Physician Assistant
- Dentist
- Emergency Medical Services
- Undecided
- Other:

What field do you want to go into? *

- General Internal medicine and hospitalist
- Internal medicine subspecialty (e.g. cardiology, nephrology, oncology)
- Pediatrics/ Pediatric subspecialty
- Ob/Gyn
- Family medicine
- Psychiatry
- Neurology
- Anesthesiology
- General surgery
- Surgical subspecialty (e.g. orthopedics, ENT, neurosurgery)
- Radiology/ Pathology
- Emergency medicine
- Undecided/I don't know yet
- Other:

What do you want to focus on? *

- Clinical / Direct patient care
- Healthcare administration
- Research, including academic & government hospitals/ research institutions
- Health-related insurance company
- Healthcare / Biotech Finance
- Healthcare industry/startup, including pharmaceuticals, medical devices / software
- Public policy
- Undecided
- Other:

Did your interest in the medicine/ healthcare profession change due to COVID-19 pandemic? *

- Increased
- Decreased
- Unchanged

Any consideration of change in career choice due to COVID-19 pandemic? *

- Yes
- No

Survey Part 5

What was your prior chosen field? *

- Undecided
- Non-Health Care Field
- Education
- Finance/ Business
- Law
- Administration
- IT/Tech/Engineering

- Physician/Medicine
- Other Healthcare Professional (e.g. nursing, physician assistant, pharmacist, dentist, etc)
- Other:

What was your reasoning for the change? *
- Personal health/ family health
- Work-life balance
- Income security
- Change of interest
- Other:

What was your previous experience with healthcare before COVID-19 pandemic? *
- Household member in medicine/ healthcare profession (people you live with e.g. parents or siblings)
- Relative/ friend in medicine/ healthcare profession (people not living with you e.g. grandparents, uncles/aunts, cousins, close family friend)
- Primary recipient of severe and/or ongoing chronic medical condition care (e.g. childhood cancer, asthma, seizure disorder)
- Household member with severe and/or ongoing medical condition care
- None of the above
- Other:

How do you consider your interaction with the American healthcare system before COVID-19 pandemic? *
- Extremely Negative 1 2 3 4 5 6 7 8 9 10 Extremely Positive

On average, how much COVID-specific media exposure have you had lately? (including, news, social media, online, etc) *
- Little to none
- Up to 1 hour/ day
- 1-2 hours/day
- 2-4 hours/day
- +4 hours/day

What is your main source of COVID-specific media/news? *
- Family/ friends/ word of mouth
- Official news outlets e.g CNN, ABC, NBC etc
- Social media
- Federal, state, and local government website (e.g. CDC, FDA, local health department)
- Other websites
- Other:

How do you feel the media coverage on the American healthcare field has been in light of COVID-19 *

(e.g. as it pertains to judging/praising healthcare professionals for their actions, judging/praising safety conditions in clinics, judging/praising quality of care, etc?)

- Extremely Negative 1 2 3 4 5 6 7 8 9 10 Extremely Positive

What are your sentiments on the future practice of medicine and hospital/clinic recovery after the pandemic? *
Financial recovery, operational recovery (e.g. enough PPE, infection control), level of government regulation/ administrative burden.

- Extremely Negative 1 2 3 4 5 6 7 8 9 10 Extremely Positive

What are your sentiments on economic and social recovery from the pandemic? *
(e.g. on unemployment returning to pre-pandemic levels, COVID-19 impacting fewer individuals health, social activities returning to normal)

- Extremely Negative 1 2 3 4 5 6 7 8 9 10 Extremely Positive

How has COVID-19 affected you personally emotionally, mentally, and/or physically on a personal level? *

- No effect 1 2 3 4 5 6 7 8 9 10 Significant effect

On average, how has COVID-19 affected your family and/or close friends on an emotional, mental, and/or physical level? *

- No effect Negative 1 2 3 4 5 6 7 8 9 10 Significant effect

Have you lost a job, grad school position, or internship as a result of COVID-19? *

- Yes
- No

Has any of your family or friends lost a job, grad school position or internship *

- Yes
- No

How has COVID-19 impacted the format of your education (check all that apply) *

- Recorded lectures/ “asynchronous lectures”
- Synchronous lectures/ “live session” only
- Online office hours (live sessions)
- Additional help through email
- No office hours/ additional help
- Other:

Within your knowledge, do you think there is increased academic dishonesty during this semester?
As a reminder, all responses are anonymous *

- 0% of class cheating 1 2 3 4 5 6 7 8 9 10 100% of class cheating

How have these changes impacted the difficulty of your courses? *

- Made classes easier
- Made classes harder

- Difficulty of classes unchanged

What contributed to less challenging courses? check all that apply *
- Open book exams
- increased time to complete exam
- Group projects in lieu of exams
- Professors more sympathetic
- No attendance/participation grades
- More generous curve
- More flexible learning environment e.g. can take breaks, re-watch lecture videos
- Other:

What contributed to more challenging courses? Check all that apply *
- Live session lectures/ office hours on inconvenient time zone
- Difficulty collaborating with group members on projects
- Disruptive/ distracting learning environment/ no dedicated study space
- Lack of technology/good internet connection
- Difficulty focusing due stress
- Additional responsibilities/ less time to study
- Other:

Has the change in academic instruction due to COVID-19 influenced your interest in medicine? *
- Increased interest
- Decreased interest
- Did not change interest
